# Fish environmental DNA in lake sediment overcomes the gap of reconstructing past fauna in lake ecosystems

**DOI:** 10.1101/2022.06.16.496507

**Authors:** Masayuki K. Sakata, Narumi Tsugeki, Michinobu Kuwae, Natsuki Ochi, Kana Hayami, Ryohei Osawa, Teppei Morimoto, Tetsu Yasashimoto, Daiki Takeshita, Hideyuki Doi, Toshifumi Minamoto

## Abstract

1. Underwater sediments are a natural archive of biological information. Reconstruction of past fauna has been conducted for various taxonomic groups using morphological remains and DNA derived from these remains. However, information on past occurrences of fish species, the top predator of lake ecosystems, could have been reproduced only in exceptional environments, and past quantitative information on fish, particularly in lake ecosystems, has been a knowledge gap in reconstructing past fauna. Tracking the quantitative fluctuations of fish is essential for reconstructing multiple trophic levels of organisms in lake ecosystems.
2. To acquire past quantitative fish information from lake sediments, we collected approximately 30 cm-length of underwater sediments in Lake Biwa. We extracted sedimentary environmental DNA (eDNA) and measured temporal fluctuations in the eDNA concentration of the native and fishery target species *Plecoglossus altivelis* and *Gymnogobius isaza*. For *P. altivelis*, we examined the possibility of tracking quantitative fluctuations by comparing sedimentary eDNA with recorded catch per unit effort (CPUE).
3. The chronology of the sediments allowed us to obtain information on sediments collected in Lake Biwa over the past 100 years. The deepest depths at which sedimentary eDNA was detected were 30 and 13 cm for *P. altivelis* and *G. isaza* from the surface, corresponding to approximately 100 and 30 years ago, respectively. In the comparison of sedimentary eDNA concentrations and biomass, we found a significant correlation between the CPUE of *P. altivelis* and its sedimentary eDNA concentration adjusted to compensate for DNA degradation. Sedimentary eDNA fluctuations were observed in *P. altivelis*, possibly reflecting the abundance fluctuation due to variations in the main food resources of zooplankton.
4. Our findings provide essential pieces for the reconstruction of past fauna of lake ecosystems. The addition of quantitative information on fish species will reach a new phase, for instance, by investigating population shifts or biological interactions in the reconstruction of past fauna in lake ecosystems.

## 1. Introduction

Lake and marine sediments are natural archives of past environmental changes and biological information. By dating the sediments and analyzing the information contained in them, an opportunity can be taken to investigate the temporal changes in ecosystems (Anderson, Renberg & Segerstrom, 1995; Davidson et al., 2011). Perspectives on past long-term changes could provide fundamental information for evaluating species evolution, responses to climate change, and human impacts, and these information could contribute to setting management and preservation strategies and understanding the factors in the establishment of ecological communities (Battarbee et al., 2005; Capo et al., 2021; Davidson et al., 2011; Ellegaard et al., 2020). Underwater sediments preserve the remains of various organisms, and lake sediments in particular have the potential to provide information related to both the underwater and terrestrial environments of the catchment area (Domaizon et al., 2017; Pedersen et al., 2015; Thomsen et al., 2012). The acquisition of past biological information using lake sediments has been conducted using morphological remains of aquatic and terrestrial macro-organisms (e.g., crustacean plankton, diatoms, pollen, plant fossils, etc.) that are well preserved in the sediments and can be visually observed and identified as proxies (Ellison, 2008; Sato et al., 2016; Frey, 1960; Hannon & Geology, 1997; Kerfoot, 1975; Leavitt & Carpenter, 1989; Prentice et al., 2018; Tsugeki, Oda & Urabe, 2003). However, challenges arise with this approach when the morphological remains are not sufficiently preserved or cannot be taxonomically identified.

To overcome these limitations, DNA derived from remains has been studied as an alternative proxy for tracking past changes in biodiversity. The use of DNA derived from remains has enabled the identification of morphologically difficult to classify remains at the species-level resolution. (Anderson-Carpenter et al., 2011; Brede et al., 2009; Coolen & Overmann, 1998; Giguet-covex & Ficetola, 2015; Härnström et al., 2011; Orsini et al., 2013; Pansu, Li et al., 2021). However, it is difficult to investigate past biological information using DNA derived from remains or morphological remains for taxa whose remains are not well preserved in sediments, such as fish (Boessenkool et al., 2014; Ellegaard et al., 2020). Because fish occupy an important ecological position as top predators in most lake ecosystems and are one of the most endangered groups of vertebrates due to habitat modification and the introduction of exotic species (Dudgeon et al., 2006; Eloranta et al., 2015), it is crucial from both the ecological and conservative perspectives to understand temporal changes in fish biomass and species composition in response to biotic and abiotic environmental changes. Therefore, the acquisition of past fish information is one of the greatest gaps in the understanding of temporal variations in lake ecosystems, and overcoming this challenge is necessary.

One of the limited ways to overcome this gap is to use sedimentary eDNA, which is the accumulation of environmental DNA (eDNA) in sediments. In the past decade, the fields of eDNA have developed and eDNA analysis is used as a powerful tool for biodiversity surveys in freshwater environments (Deiner et al., 2017; Goldberg et al., 2016; Taberlet et al., 2012; Thomsen et al., 2012). Generally, eDNA contained in water is used for current biodiversity monitoring (Blabolil et al., 2021; Miya, 2022; Riaz et al., 2020; Sakata et al., 2021), and is degraded quickly in water (Barnes & Turner, 2016; Stewart, 2019; Jo, Takao & Minamoto, 2021). By contrast, sedimentary eDNA is barely degraded and can be detected for a long time (Ogata et al., 2021; Sakata et al., 2020; Turner, Uy & Everhart, 2015). Therefore, sedimentary eDNA from past sediments provides biological information on past fish. Previous studies have successfully detected fish eDNA from past sediments in lakes and seas (Kuwae et al., 2020; Nelson-Chorney et al., 2019; Stager et al., 2015). For example, Nelson-Chorney et al. (2019) showed that previous non-native fish invasion was detected by eDNA, and the estimated date of the sediment layer where eDNA was detected corresponded with the records of introduction. Kuwae et al. (2020) tracked changes in fish abundance based on the fluctuation of sedimentary eDNA concentrations in the sea. However, these previous studies were conducted in limited areas suitable for long-term preservation of DNA, such as lakes at high altitudes and high latitudes with low-temperature environments or sea areas under anoxic conditions, and it has been suggested that DNA is difficult to detect in aerobic environments (Kuwae et al., 2020). By contrast, there are no reports of sedimentary eDNA detected from past sediments in lakes in temperate zones, where eDNA degradation seems to be relatively rapid. Because biodiversity is richer in temperate zones than in high-latitude zones (Fine, 2015; Hillebrand, 2004), temperate zones are also important regions from the perspective of biodiversity conservation. Demonstrating the feasibility of past reconstruction of fish fauna and their quantitative biomass in temperate zones will play a valuable role in filling gaps in our knowledge of biological states and tracking ecosystem changes. In particular, quantitative reconstruction of fish in lake ecosystems will be the last step needed to reconstruct the biological information of the entire lake ecosystem.

In this study, we collected sediment cores from Lake Biwa, the largest lake in Japan, and attempted to detect fish using sedimentary eDNA to investigate whether fish information can be recovered from past lake sediments in a temperate zone. Specifically, we aimed to (1) demonstrate whether fish sedimentary eDNA can be detected even in a temperate environment, (2) examine whether the obtained eDNA signals reflect biomass fluctuations, and (3) explore the environmental factors that cause fish fluctuations in the past. These findings are essential for our understanding of lake ecosystem changes.

## 2. Materials and Methods

### 2.1. Target species and real-time PCR assays

We set two fish species, Plecoglossus altivelis altivelis and Gymnogobius isaza, as the target species. Plecoglossus altivelis is an important fishery resource, and catch per unit effort (CPUE) was recorded between 1970 and 2010 in Lake Biwa (Liu et al., 2020; Suzuki & Kitahara, 1996). Gymnogobius isaza is endemic to Lake Biwa and is also a fishery resource (Nagoshi, 1966). In Lake Biwa, these two species are advantageous due to as it is possible to evaluate the validity of their detection based on records of their catches. Real-time PCR assay for *P. altivelis* was performed as previously described (Yamanaka & Minamoto, 2016). However, there has been no eDNA study for *G. isaza*, therefore, we developed a real-time PCR assay for *G. isaza*.

Sequences of the mitochondrial cytochrome b (cytb) gene of the target species *G. isaza* and the closely related species, *G. urotaenia* a potential sympatric species in Japan, were downloaded from the database of the National Center for Biotechnology Information (NCBI: https://www.ncbi.nlm.nih.gov). Based on these sequences, species-specific primers satisfying three conditions were designed according to previous studies: 1) melting temperature around 60 °C; 2) at least two specific bases within the five bases at the 3’ ends of both forward and reverse primers; and 3) targeting a small fragment (designed for an amplicon size of 50–150 bp) (Goldberg et al., 2016; Sakata et al., 2017; The eDNA Society, 2019). The sequences used to design the primers are listed in Table 1. The designed primers and probes are listed in Table 2. The amplicon length was 88 base pairs (bp). In addition, potential cross-reactivity of the assay was checked by in silico analysis (i.e., Primer-BLAST was performed on all databases; https://www.ncbi.nlm.nih.gov/tools/primer-blast/index.cgi?LINK_LOC=BlastHome).

**Table 1.** Accession numbers of the nucleotide sequences used for designing species-specific primers and a probe for *Gymnogobius isaza*.

| Species name | Accession Nos. |  |  |
| --- | --- | --- | --- |
| *Gymnogobius isaza* | LC098530.1, | LC098531.1, | LC098529.1, |
|  | LC098528.1 |  |  |
| *Gymnogobius urotaenia* | KT601093.1, | LC098543.1, | LC098545.1, |
|  | LC098542.1 |  |  |

**Table 2.**
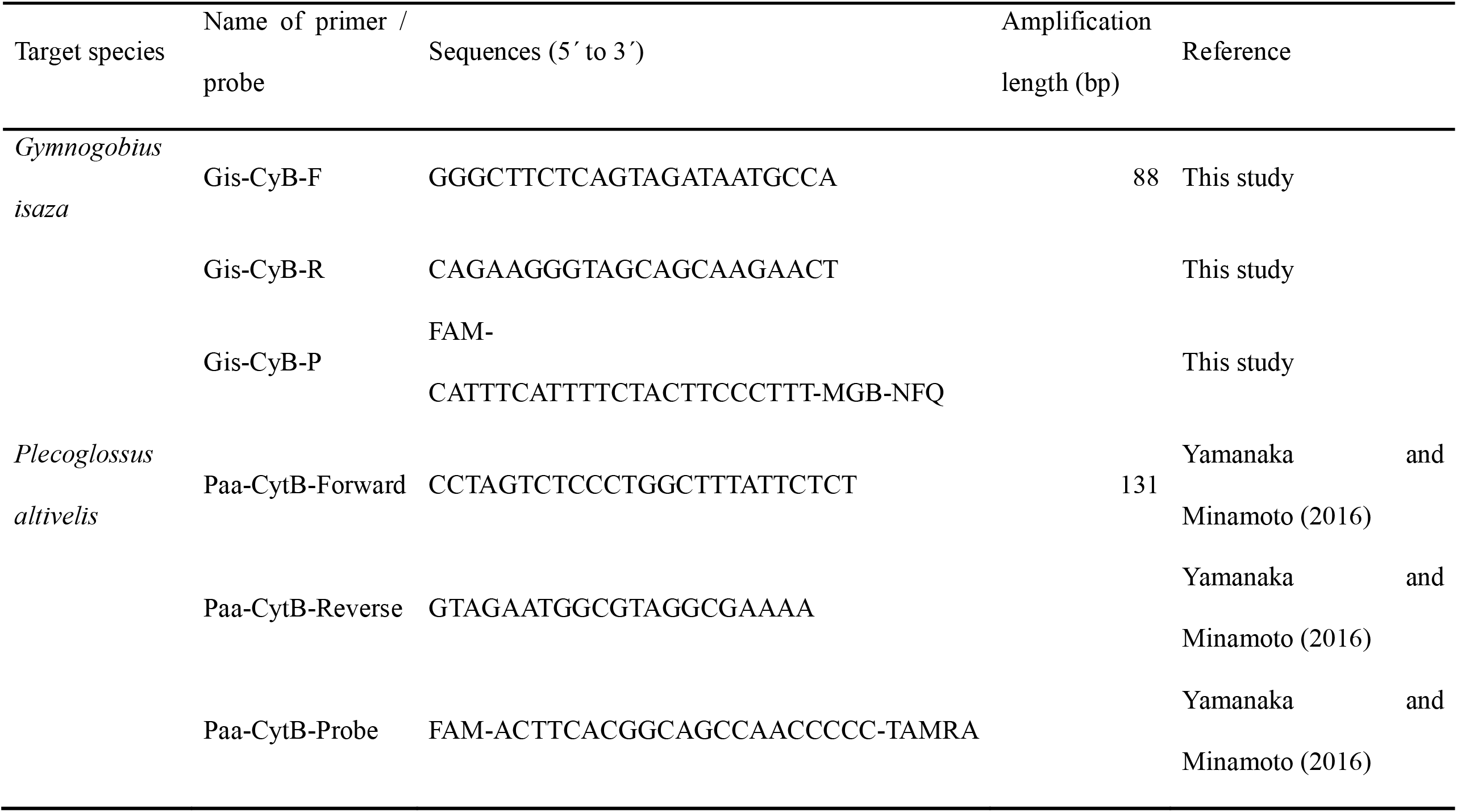
Primers and probes used in qPCR experiments.

The specificity of the assay was evaluated by real-time PCR using target and nontarget DNA templates. Total DNA was extracted from the tissues of the target and related species using the DNeasy Blood & Tissue Kit (QIAGEN Science, Hilden, Germany) following the manufacturer’s instructions. The concentration of DNA was measured using a fluorescence quantification apparatus (Qubit 3.0 Fluorometer; ThermoFisher Scientific, Massachusetts, USA). Real-time PCRs were carried out in triplicate using extracted DNA from each species as a template with a CFX96 Real-Time System (Bio-Rad, Hercules, CA, USA). Each reaction (20-μL final volume) contained 900-nM primers, 125-nM TaqMan probe, 0.1 μL of AmpErase Uracil N-Glycosylase (ThermoFisher Scientific, Massachusetts, USA), and 100 pg DNA template of each species in 1× Environmental Master Mix 2.0 (ThermoFisher Scientific, Massachusetts, USA). The real-time PCR conditions were as follows: 2 min at 50 °C, 10 min at 95 °C, and 55 2-step cycles of 15 s at 95 °C, and 60 s at 60 °C. To detect false positives due to contamination during the real-time PCR procedures, ultrapure water was used instead of DNA in the three reaction mixtures (non-template negative controls).

The limit of detection (LOD) and limit of quantification (LOQ) were measured for each assay. The concentration of the standards was adjusted to 3,000, 300, 30, 15, 10, 5, 3, and 1 copy per reaction. For each assay, PCR was performed in triplicate using standard and negative controls. The PCR conditions were the same as described above. The LOD and LOQ were the lowest eDNA concentrations (copies per reaction) at which DNA amplification was observed in one of the triplicates and in all of the triplicates, respectively (Agersnap et al., 2017; Harper et al., 2019; Takeshita et al., 2020).

### 2.2. Sediment core sampling for reconstructing past biological information

To reconstruct the past fish information using sedimentary eDNA from bulk sediment, sediment core sample (ADG7 core) was collected from the north basin of Lake Biwa at a pelagic site (water depth of 71.5m; 35°15’10.1’’N, 136°03’48.7’’E) with a 10.9-cm-inside-diameter gravity corer on August 27, 2019 (Fig. 1). The core tube was cleaned with 0.6% sodium hypochlorite, tap water, and Milli-Q water before use. The core was carefully cut in 1-cm increments from the surface to a depth of 31 cm. Each increment was preserved in a shield bag and stored at –25°C until DNA extraction. Knifes and cutting apparatus were cleaned with 0.6% sodium hypochlorite, tap water, and Milli-Q water before use. To check for DNA contamination between the samples, the instruments were rinsed with approximately 100 mL of distilled water every 15 slices, and water was used as a negative control.

**Fig. 1.**
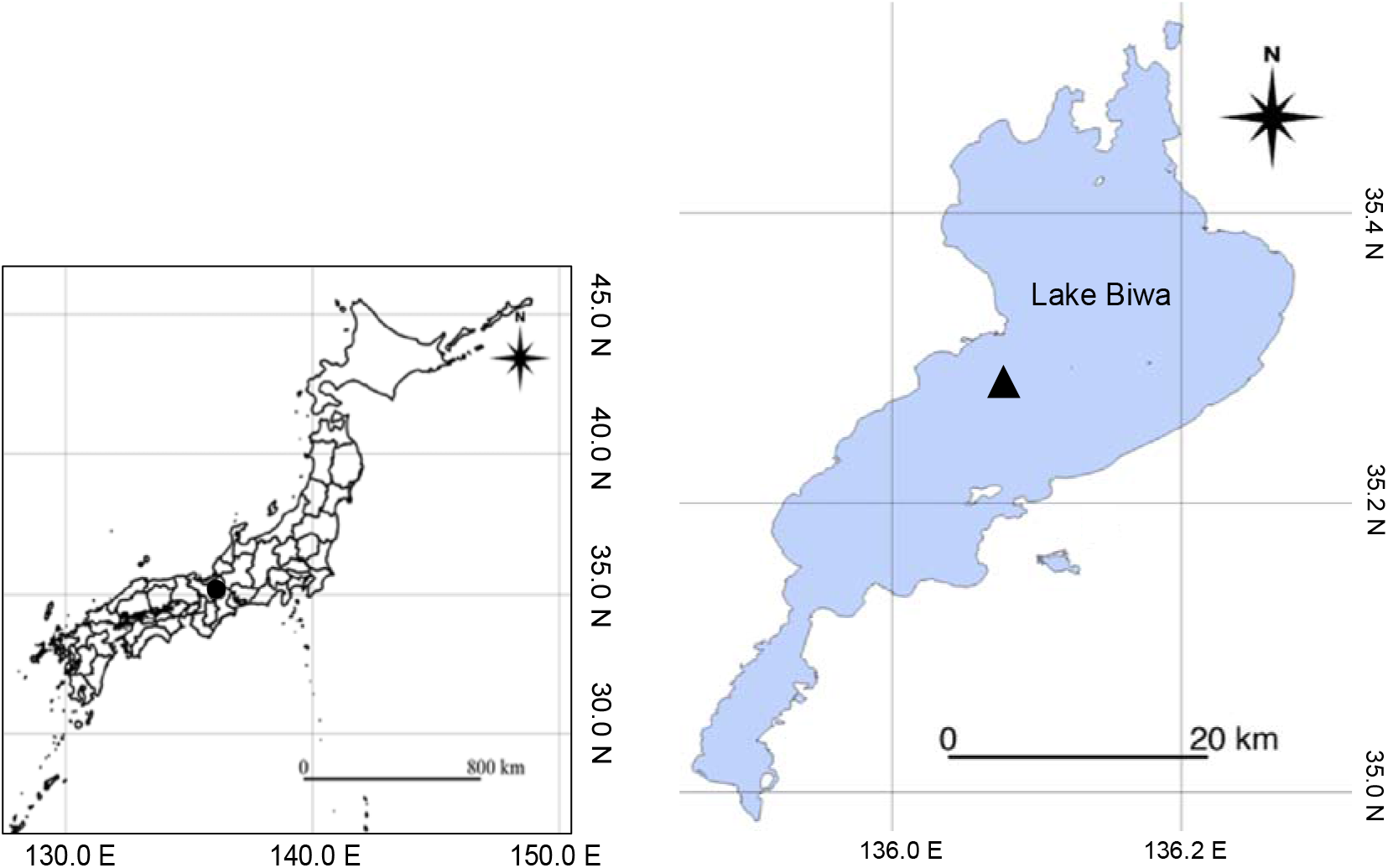
Location of the sampling site. The sampling site is located offshore in the northern area of Lake Biwa (water depth of 71.5m; 35°15 N, 136°03 E). The circle in the left panel indicates the location of Lake Biwa. The triangle in the right panel indicates the sampling site. Maps are based on the Digital Map (Basic Geospatial Information) published by the Geospatial Information Authority of Japan (http://maps.gsi.go.jp/)

### 2.3. Dating and measurement of chlorophyll concentration

The chronological age of the ADG7 core was estimated indirectly by comparison with the profiles of chlorophyll pigments and magnetic susceptibilities of another chronological LB7 core collected at the same point in this study and previously performed sediment chronology analysis by Tsugeki et al. (2021). The chronology of the LB7 core was determined based on the constant rate of supply (CRS) method of 210Pb dating (Appleby & Oldfield, 1978) and verified using the 137Cs peak traced from 1962 to 1963 (Appleby, 2001).

The chronology of ADG7 was determined by comparing the reference layers of chlorophyll pigments and magnetic susceptibilities of the ADG7 and LB7 cores. In the ADG7 and LB7 cores, magnetic susceptibility and chlorophyll pigments were measured using an SM-30 m (ZH instruments, Brno, Czech Republic) and a UV–Vis mini 1240 spectrophotometer (Shimadzu, Kyoto, Japan), respectively. A detailed method for determining the concentrations of chlorophyll pigments (chlorophyll-a and phaeopigments) has been reported by Tsugeki et al. (2021). Briefly, chlorophyll pigments in sediment samples (approximately 1.0 g wet weight) were extracted in 10 mL of 90% acetone at room temperature. The samples were then centrifuged, the chlorophyll-a and its fractions in the supernatant were measured using a spectrophotometer, and their concentrations were calculated according to the method of Lorenzen (1967).

In the profile of chlorophyll pigments and magnetic susceptibilities, layers showing distinctive peaks and troughs were used as reference layers. Thus, the age of the reference layer of the ADG7 core was determined by comparing it with the dated LB7 core reference layer, and the chronology of the ADG7 core was determined based on its depositional rate of the ADG7 core.

### 2.4. Sedimentary eDNA extraction

Sediment samples were mixed well and subsequently separated into three subsamples (10 g each) from a single sediment slice. Sedimentary eDNA was extracted from 10 g sediment samples by combining alkaline DNA extraction (Kouduka et al., 2012) with ethanol precipitation and a fecal/soil DNA extraction kit (PowerSoil DNA Isolation Kit; Qiagen), according to the methods of a previous study with minor modifications (Sakata et al., 2021). A DNA enhancer (G2 DNA/RNA Enhancer; Ampliqon, Odense, Denmark) was added during extraction (Jacobsen et al., 2018). To detect cross-contamination during the extraction process, 10 mL of ultrapure water was used as an extraction negative control and treated in the same way as the sediment samples. The final volume of the eDNA was 100 μL. All tools were either disposable consumables or decontaminated with chlorine bleach (0.1% effective chlorine concentration).

### 2.5. Quantification of fish sedimentary eDNA

The eDNA concentrations of *P. altivelis* and *G. isaza* in each eDNA sample were quantified by TaqMan real-time quantitative PCR (qPCR) targeting the cytochrome b region of *P. altivelis* and *G. isaza* using previously developed assays for *P. altivelis* (Yamanaka & Minamoto, 2016) and assays for *G. isaza* developed in this study. Primers and probes used in this study are listed in Table 2. qPCR was performed in triplicate using extracted eDNA from each sample as a template. Each reaction (20-μL final volume) contained 900-nM primers, 125-nM TaqMan probe and, 0.1 μL of AmpErase Uracil N-Glycosylase (Thermo Fisher Scientific) in 1× Environmental Master Mix 2.0 (Life Technologies) and 2 μL eDNA. The real-time PCR conditions were as follows: 2 min at 50 °C, 10 min at 95 °C, and 55 cycles of 15 s at 95 °C, and 60 s at 60 °C. To obtain calibration curves, a dilution series of standards (3 × 101 to 3 × 104 copies in each reaction) was simultaneously quantified. The standards were linearized plasmids containing the synthesized artificial DNA fragments of the target cytb gene sequence of *P. altivelis* and *G. isaza*. Ultrapure water was used instead of DNA in the three reaction mixtures as a non-template negative control. The DNA concentration in each sample was calculated as the average of the three qPCR replicates. When a negative detection was obtained in any of the replicates, the DNA concentration of that replicate was assigned zero (Ellison et al., 2006).

To check for PCR inhibition, all eDNA samples were spiked with 2,000 copies of lambda phage DNA as an internal positive control (IPC), and qPCR was conducted targeting lambda phage DNA. This test was carried out using the Lambda-7184F (5-TTCTCTGTGGAGGAGTCCATGAC-3) and Lambda-7267R (5-GCTGACATCACGGTTCAGTTGT-3) primers and the TaqMan probe Lambda-7210P (5-FAM-AGATGAACTGATTGCCCGTCTCCGCT-TAMRA-3) (Honjo et al., 2010). The results of the inhibition test showed no inhibition for all samples (ΔCt (Ct sample - Ct positive control) < 0.1, corresponding to a difference of < 1.07 times between the estimated eDNA copy numbers).

### 2.6. Statistical analysis

Because the total chlorophyll a and its derivative concentrations (Fig. 2a) may have decreased over time due to biotic and abiotic degradation in sediments (Leavitt & Carpenter, 1990; Ming-Yi, Lee & Aller, 1993), the total chlorophyll a and its derivative concentrations in the older sediment layers would have been underestimated. Therefore, in accordance with a previous study, we estimated their concentrations before biotic and abiotic degradation. In a previous study, the concentrations of total chlorophyll a and its derivatives before degradation after burial were reconstructed by calculating the degradation rate along first-order kinetics assuming pigment degradation (Tsugeki et al., 2017). Here, we used the following equation to calculate the degradation rate, according to a previous study:

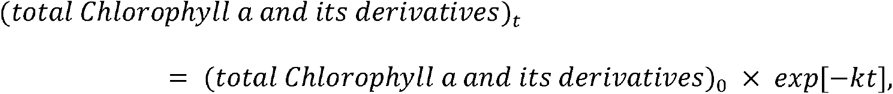

where (total chlorophyll a and its derivatives)t is the concentration of total chlorophyll a and its derivatives at time t, t is the time point after sedimentation of total chlorophyll a and its derivatives (=0 at the sediment surface), and k is a first-order rate constant (year-1). Based on this equation, the following equation was obtained from the data:

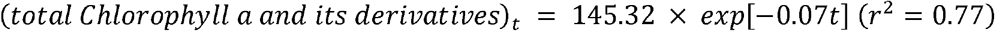

**Fig. 2.**
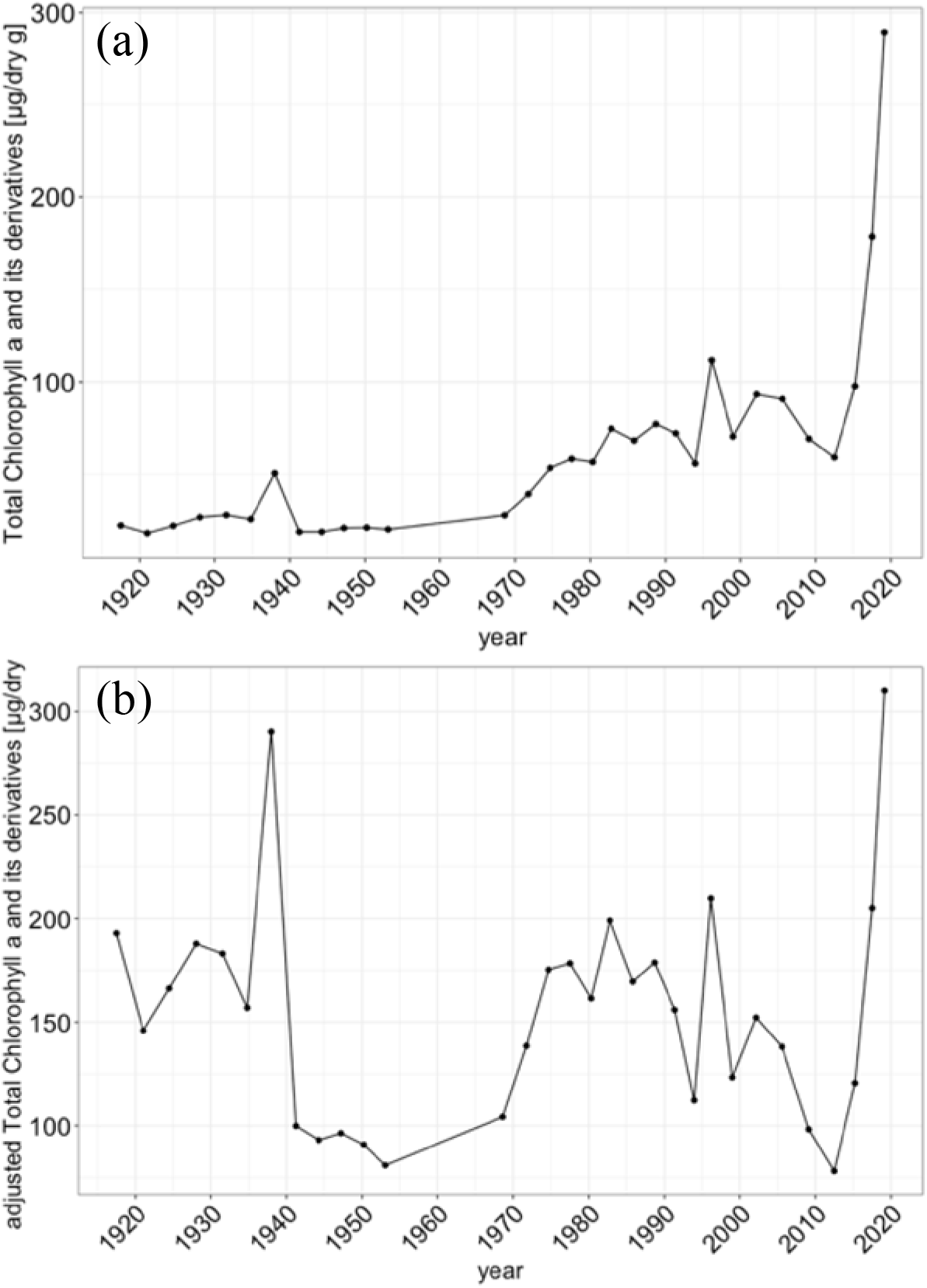
Time profiles of concentrations of total chlorophyll a and its derivatives (chlorophyll-a + phaeopigments) dated by inter-core comparison. The points indicate the measured total chlorophyll concentration (chlorophyll-a and phaeopigments concentration). (a) The concentrations of total chlorophyll a and its derivatives is not adjusted for degradation. (b) The concentrations of total chlorophyll a and its derivatives concentration is adjusted for degradation.

Based on this equation, the concentration before degradation was estimated using the following equation.

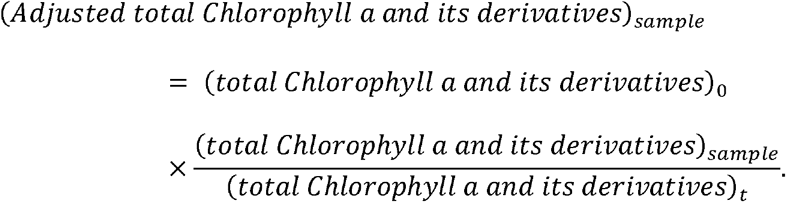

Biotic or abiotic degradation is also expected in eDNA (Sakata et al., 2020; Wei, Nakajima & Tobino, 2018;); however, the detection rate is relatively low and difficult to model. Therefore, because there was a correlation between chlorophyll pigments (not adjusted) and eDNA (linear model, p < 0.05; Fig. S1, S2), the concentration of eDNA was adjusted using the coefficients of the model equation for chlorophyll pigments, using the following equation:

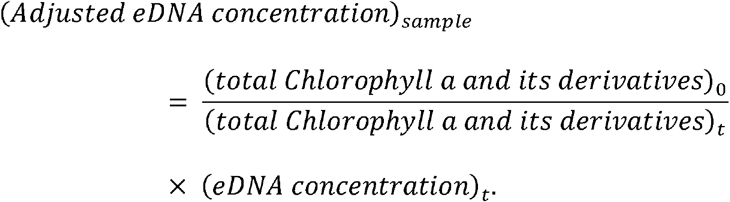

We evaluated the relationship between sedimentary eDNA concentrations and biomass using a generalized linear model (GLM) with negative binomial distribution using the function glm.nb in R package MASS. A negative binomial regression with a log-link function was used to account for overdispersion (Ver Hoef & Boveng, 2007). We used “eDNA concentration (not adjusted)” or “adjusted eDNA concentration” as a dependent variable, and “CPUE” as the independent variable. This analysis was performed only for *P. altivelis* because the number of detections of *G. isaza* was low and could not be compared. The CPUE of *P. altivelis* calculated in a previous study was used as an indicator of the *P. altivelis* biomass (Liu et al., 2020). In this analysis, the average CPUE for four years included by each core slice based on the top and bottom years of each slice was used as in Kuwae et al. (2020). All analyses were performed using R ver. 3.6.3 (R Core Team, 2020).

## 3. Results

### 3.1. Validation of developed real-time PCR assays

The specificity of the primers designed for *G. isaza* was confirmed using in sillico testing. In addition, real-time PCR with 100 pg of the total DNA of the most related species (G. urotaenia) as a template showed no amplification in any of the three replicates. Thus, the real-time PCR assay designed in this study had sufficient specificity for detecting target species in our survey area. For the *P. altivelis* assay, the LOQ and LOD were five copies and three copies, respectively. For the *G. isaza* assay, both LOQ and LOD were one copy.

### 3.2. Chronology of the ADG7 core

Profiles for magnetic susceptibilities and chlorophyll pigments were compared between the ADG7 core and LB7 core to estimate the chronology of ADG7 (Fig. S3 and S4). Based on a comparison of the chlorophyll pigments,, two depths were used as reference layers for the pigment proxies, which increased at a depth of 21.5 cm layer (each layer was expressed as mid-depth; 21.5 cm for the 21–22 cm depth layer) in core LB7 (estimated date: 1953.1 ± 12.1), peaked at a depth of 13.5 cm (1982.8 ± 2.5), whose peak were comparable to those of the 19.5 cm and 13.5 cm layers in ADG7 (Fig. S3). Similarly, in comparison of magnetic susceptibilities, two reference layers were detected (Fig. S4). The first reference layer of the magnetic susceptibilities peak was at the 9.5 cm depth layer in LB7 (estimated date: 1996.2 ± 0.7) comparable to that of the 8.5 cm depth layer in ADG7. The second reference layer was at a depth of 17.5 cm in LB7 (estimated date: 1968.6 ± 5.9) and was comparable to that at 18.5 cm in ADG7. Based on the ages of these four reference layers in the LB7 core, the chronology of the ADG7 core was determined. The oldest layer (31 cm bottom depth) in the ADG7 core was estimated to be 1917. The chlorophyll pigments profile adjusted for degradation is shown in Fig. 2b.

### 3.3. Sedimentary eDNA analysis

The R2 values of the calibration curves were >0.98 in all real-time PCR runs for sedimentary eDNA analysis. The values of slopes, intercepts, and PCR efficiency were, respectively, −3.609 to −3.454, 41.894 to 42.943, and 89.3% to 101.1% for *P. altivelis*, −3.347 to −3.200, 39.666 to 40.056, and 99.0% to 105.4% for *G. isaza.* For *P. altivelis*, sedimentary eDNA was detected in 17 subsamples from 13 layers of 93 subsamples from 31 slices (Fig. 3a). For *G. isaza*, sedimentary eDNA was detected in 14 subsamples from seven out of 93 subsamples from 31 sliced layers (Fig. 3b). The deepest depths at which sedimentary eDNA was detected were at 29.5 cm (estimated date: before 1924) and 12.5 cm depth (1985.8) layers from the surface for *P. altivelis* and *G. isaza* (Fig. 3a and 3b), corresponding to approximately 100 and 30 years ago, respectively. None of the negative controls (negative controls, extraction negative controls, and PCR negative controls) showed amplification. eDNA fluctuations adjusted for degradation in P. altiveli and *G. isaza* are shown in Fig. 3c and 3d, respectively. No significant relationship was observed between CPUE (Fig. S5) and non-adjusted sedimentary eDNA concentration of *P. altivelis* (p = 0.59; Fig. 4a). In contrast, a significant relationship was found between CPUE and adjusted sedimentary eDNA concentration of *P. altivelis* (p < 0.05; Fig 4b).

**Fig. 3.**
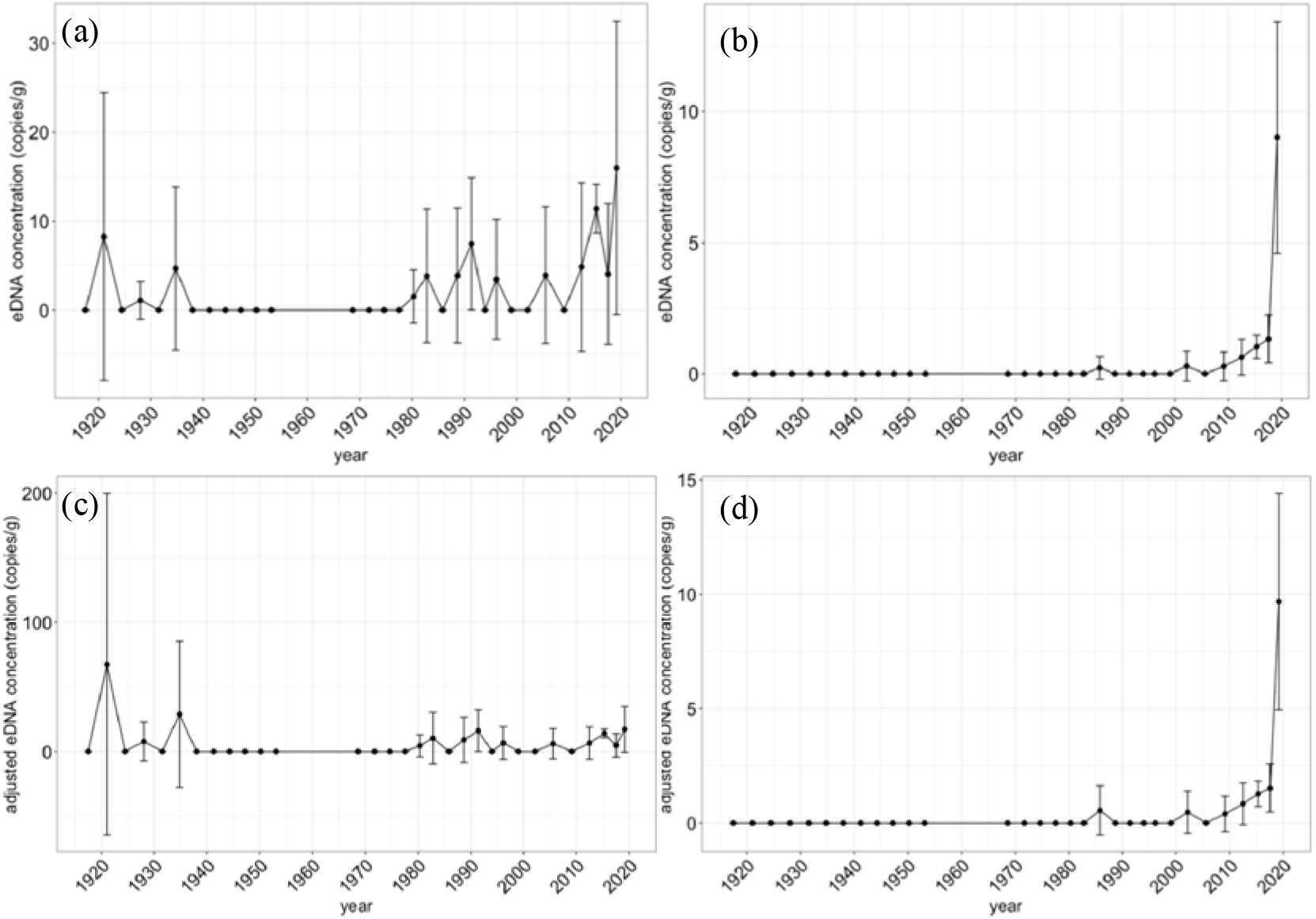
Time profiles of eDNA concentrations of target species dated by inter-core comparison. Points and error bars indicate the mean and standard deviation of sedimentary eDNA concentration from the three subsamples, respectively. (a) The profile of *Plecoglossus altivelis* is shown, and eDNA concentration is not adjusted for degradation. (b) The profile of *Gymnogobius isaza* is shown, and eDNA concentration is not adjusted for degradation. (c) The profile of *Plecoglossus altivelis* is shown, and eDNA concentration is adjusted for degradation (see text for details). (d) The profile of *Gymnogobius isaza* is shown, and eDNA concentration is adjusted for degradation.

**Fig. 4.**
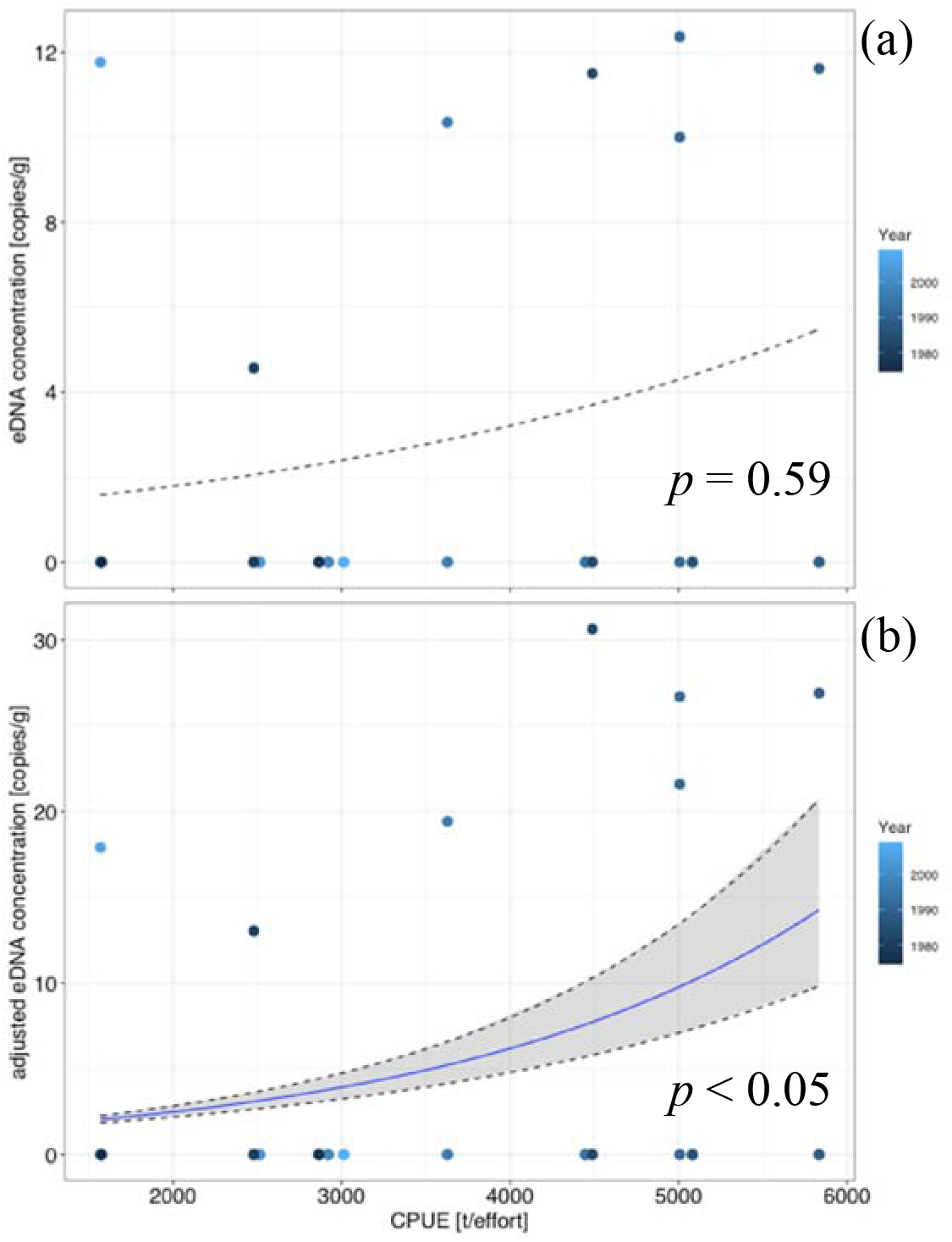
(a) Relationships between eDNA concentrations and the recorded CPUE for *Plecoglossus altivelis.* The regression model showed no significant relationship between the variables. The dashed line indicates the estimated regression line. The color of the dots indicates the age of the estimated sediment layer. (b) Relationships between adjusted eDNA concentrations and the recorded CPUE for *Plecoglossus altivelis.* The regression model showed that there was a significant positive relationship between the variables. The solid line indicates the estimated regression line and shaded area indicates 95% CI. The color of the dots indicates the age of the estimated sediment layer.

## 4. Discussion

The detection of fish sedimentary eDNA, other than in special environments where DNA is easily conserved, as in previous studies (Kuwae et al., 2020; Nelson-Chorney et al., 2019), is an advance in the reconstruction of lake ecosystems; the detection of fish sedimentary eDNA will overcome the gap in reconstructing past fauna in lake ecosystems. To achieve this, we attempted to track the fluctuations of *P. altivelis* and *G. isaza*, which are fishery resources, using sedimentary eDNA from temperate Lake Biwa. Sedimentary eDNA from 100 and 30 years ago has been successfully detected in *P. altivelis* and *G. isaza*, respectively. In particular, for *P. altivelis*, the adjusted sedimentary eDNA, considering the degradation effect, correlated with historical CPUE data. In environments where eDNA is relatively easily degraded, such as temperate environments, considering the degradation effect may provide more robust results.

In this study, fish sedimentary eDNA dating back to the past 100 years was detected in a temperate lake where DNA is relatively easily degraded. This may have been achieved by using sample volumes that were three times larger than those used in previous studies (Kuwae et al., 2020; Nelson-Chorney et al., 2019). The use of an alkaline buffer prior to purification with commercial kits allows handling of large sample volumes, regardless of the volume limitations of the commercial kit (Sakata et al., 2020). Even if eDNA is heterogeneously distributed or present in low amounts in the sediment (Sakata et al., 2021), it would be possible to detect it using a larger sample volume. Although only a few positive signals of eDNA for *G. isaza* was detected in this study, when compared with the CPUE variation in *G. isaza* (1962–2002) (Briones et al., 2012), eDNA signals were detected in the sediment layer corresponding to the peak of the CPUE around 1980. This suggests that eDNA can be detected in *G. isaza* in association with biomass variations. Recent detection of eDNA may also suggest that the biomass of *G. isaza* is increasing. Although a simple comparison is difficult owing to the small number of detections, improving detection sensitivity would be one of the solutions to this problem (see below for details).

We showed that fluctuations in the adjusted eDNA concentration were correlated with the CPUE of *P. altivelis.* Thus, it would be possible to reconstruct past biomass fluctuations by detecting temporal eDNA fluctuations in sediment cores. In addition, in *G. isaza*, as in *P. altivelis*, the fluctuations in eDNA adjusted for degradation would reflect fluctuations in biomass. This fluctuation in *P. altivelis* biomass may be influenced by fluctuations in organisms with lower trophic levels than fish, represented by zooplankton. The main food resource of *P. altivelis* in Lake Biwa is zooplankton of the genus Daphnia, and *P. altivelis* are known to selectively prey on these organisms (Kawabata et al., 2002; Kawabata, Narita & Nishino, 2006). During the 1920s and 1940s, *P. altivelis* eDNA was relatively stable, but during the 1940s–1970s, almost no eDNA was detected. This period coincided with the period when many resting eggs of *Daphnia* were detected (Tsugeki et al., 2022). *Daphnia* and other zooplankton species produce resting eggs because of habitat deterioration or stress (Hairston, 1996). In Lake Biwa, the water level dropped from the 1940s to 1970s due to reclamation, and the environment would have changed drastically (Nishino, 2020). In fact, the concentration of chlorophyll, adjusted for degradation, declined sharply in 1940, suggesting a major environmental change. Since in Lake Biwa, *Daphnia* were kept low abundance before the 1960s (Tsugeki *et al.*, 2022), thus it is likely that highly limited food resources during the 1940s~1970s had been keeping the *P. altivelis* population low abundance, resulting to become undetectable of eDNA due to a low amount of eDNA. In addition, since eDNA of *Eodiaptomus japonicus*, one of the major food resources of *P. altivelis*, tends to be detected at relatively slightly higher concentrations before 1940 (Liu et al. 2021; Nakane et al. unpublished), the abundance of this food resource of *P. altivelis* may have this abundance may have contributed to the high biomass of *P. altivelis* during this period. *P. altivelis* eDNA concentrations have been increasing since the 1980s when *Daphnia* population were kept high abundance (Tsugeki *et al.*, 2022). The zooplankton increases like *Daphnia* as food resources were assumed to provide an increase in the biomass of *P. altivelis* which resulted in an increase in eDNA concentrations. This increase in zooplankton is estimated to have been influenced by eutrophication associated with anthropogenic activities since the 1980s, and the eutrophication of the lake increases nutrient concentrations, which in turn increases phytoplankton, which are primary producers (Tsugeki et al., 2010). This causes an increase in the abundance of lower consumers, such as zooplankton, which in turn increases the biomass of fish, the top predator of the lake ecosystem. This may indicate temporal fluctuations in the ecosystem due to bottom-up effects (McQueen, Post & Mills, 1986). The community dynamics caused by such bottom-up effects are compatible with the theory that these effects dominate community dynamics in eutrophic environments, such as Lake Biwa, during periods of increasing *P. altivelis* (McQueen et al., 1986).

Reconstructing past fluctuations of *P. altivelis* and *G. isaza* in Lake Biwa will contribute to the understanding of how fish populations change due to environmental changes. If the response of fish populations to environmental change is known, it may be possible to predict future directions by building predictive models for future environmental changes. This is also possible in other fish species. These findings are also important for the resource management of these useful fisheries species. In addition, since quantitative reconstructions are possible, past reconstructions of multiple species with various ecological characteristics may also provide insight into changes in biotic interactions if variations in the biomass of multiple species over time are obtained, as shown in a previous study (Ushio et al., 2018). If the past dynamics of fish are provided by sedimentary eDNA, information on higher-order consumers, which has been the gap of past reconstruction in lake ecosystems (Ellegaard et al., 2020), will be obtained by analyzing these quantitative data using time series analysis. It may become possible to reconstruct the dynamics of the overall lake ecosystem and observe the application of the ecosystem to environmental changes. Such observations will contribute to the understanding of the mechanisms that drive ecosystems and environmental factors that affect communities.

The difference in the oldest estimated age at which eDNA was detected in sediments between the two target species and the intermittent detection of the two species were issues identified in this study. Because the LOD experiments showed sufficient detectability of the detection assays, these are assumed to be due to factors including the initial concentration of eDNA and spatial variation in sedimentary eDNA within the study area (Sakata et al., 2021). As the concentration of aqueous eDNA is affected by biomass (Doi et al., 2017; Salter et al., 2019; Takahara et al., 2012), the initial concentration of sedimentary eDNA should also be affected by biomass. Low initial concentrations of eDNA accumulated in the sediment can reduce the detectable time of eDNA in the sediment owing to degradation over time. In addition, because the shedding rate of eDNA differs depending on the fish species, the amount of eDNA released between the two target species may have differed (Andruszkiewicz et al. 2020), affecting the initial concentration of sedimentary eDNA. Enhanced detectability will contribute to improving these issues by increasing the number of sampling points, number of cores, sample volumes, and extraction efficiency. In addition, abiotic factors such as water quality and water temperature are likely to influence the accumulation and persistence of sedimentary eDNA. However, because the detection of sedimentary eDNA and its relationship to environmental factors is still unknown, the relationship between these factors should also be investigated in future studies.

In this study, we successfully detected fish DNA from a sediment core in a temperate lake and showed that fluctuations were correlated with changes in biomass. Tracking fish fluctuations in the past has overcome the gap in the reconstruction of fauna in lake ecosystems. In addition, comparing data from previous historical reconstructions of low-order consumers with historical reconstructions of fish, which are high-order predators in lake ecosystems, will allow for discussion, along with a wide range of trophic levels. This will make it possible to reconstruct the biota shifts in the entire lake ecosystem from sediments, which may provide clues to the response of the ecosystem to environmental change and to unravel the drivers of the ecosystem.

## Supporting information

Supporting Infomation

## Acknowledgements

We are grateful to M. Honjo, S. Goda, and T. Akatsuka for their assistance with the laboratory analysis and field sampling. This study was supported by Grants-in-Aid for Scientific Research (Nos. 17K00528 and 22K15183) from the JSPS. This study was supported by a Grant-in-Aid for JSPS Research Fellows (no. JP19J11126). This study was supported by the Joint Usage/Research Grant of the Center for Ecological Research, Kyoto University (2017-2019 jurc-cer), and the Academic Research Organization Joint Usage/Research Grants from Leading Academia in Marine and Environment Pollution Research (LaMer), Ehime University.

## Author Contributions

MKS, NT, MK and TM conceived and designed the study. MKS, NT, MK, NO, KH, RO, TM, TY and DT performed the field sampling and subsequent processes. MKS, NT, MK, NO and KH conducted the laboratory analysis. MKS, NT, HD and TM contributed to the statistical analysis. MKS and TM wrote and edited the first draft of the manuscript. All authors discussed the results and contributed to the development of the manuscript.

## Conflict of Interests Statement

The authors declare that they have no known competing financial interests or personal relationships that could have influenced the work reported in this study.

## Data Availability Statement

The raw data will be deposited to Dryad upon acceptance.

## Notes

### Competing Interest Statement

The authors have declared no competing interest.

