## Supporting Infomation for "Fish environmental DNA in lake sediment overcomes the gap of reconstructing past fauna in lake ecosystems"

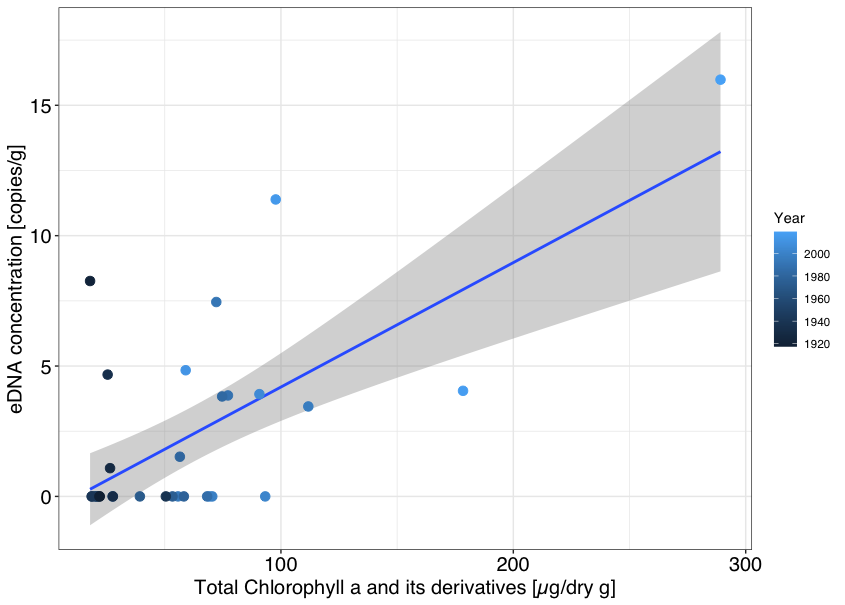

Fig. S1. Relationships between *Plecoglossus altivelis* eDNA concentrations and concentrations of total chlorophyll a and its derivatives. The blue line and the gray area indicate the regression line and 95% confidence intervals, respectively. The color of the dots indicates the age of the estimated sediment layer. The regression model showed the significant positive correlation between the sedimentary eDNA concentration and concentrations of total Chlorophyll a and its derivatives.

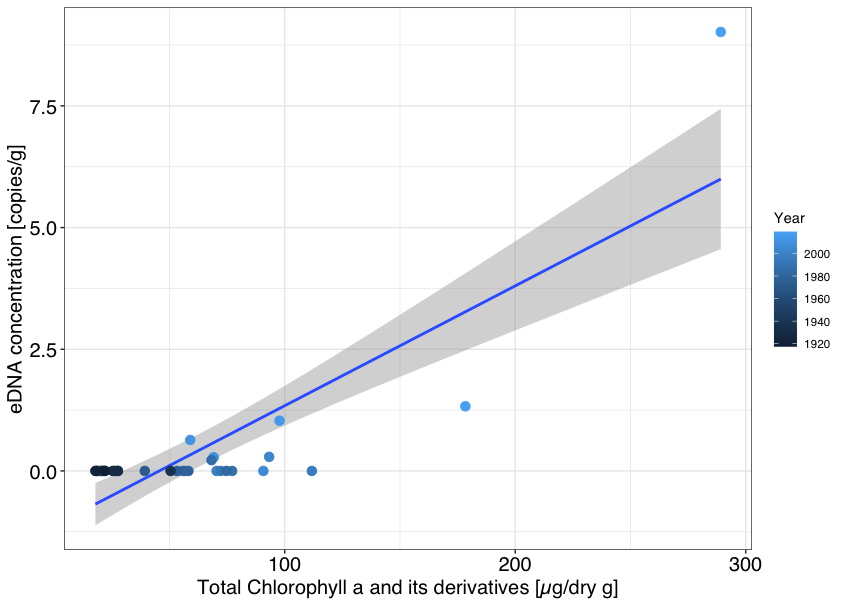

Fig. S2. Relationships between *Gymnogobius isaza* eDNA concentrations and concentrations of total Chlorophyll a and its derivatives. The blue line and the gray area indicate the regression line and 95% confidence intervals, respectively. The color of the dots indicates the age of the estimated sediment layer. The regression model showed the significant positive correlation between the sedimentary eDNA concentration and concentrations of total Chlorophyll a and its derivatives.

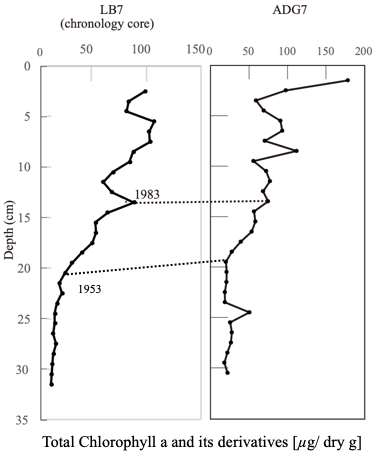

Fig. S3. Comparison of the proxy of concentrations of total chlorophyll a and its derivatives. The ADG7 core was sampled in this study. The LB7 core was sampled in a previous study and dated using the constant rate of supply (CRS) model of ^210^Pb dating.

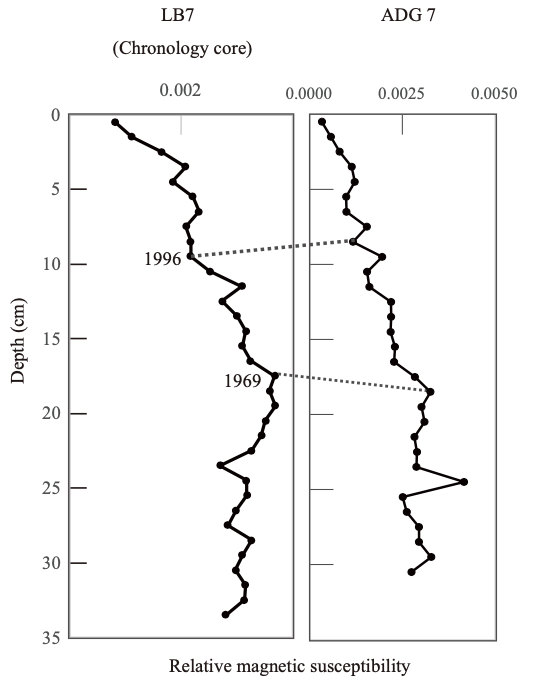

Fig. S4. The comparison of proxy of magnetic susceptibility. The ADG7 core was sampled in this study. The LB7 core was sampled in a previous study and dated using the CRS model.

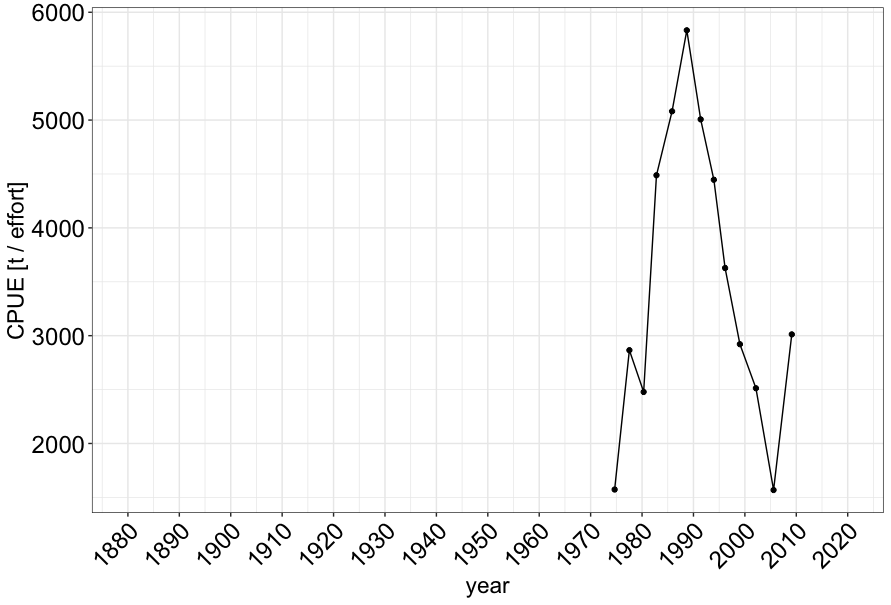

Fig. S5. The variation of CPUE for *Plecoglossus altivelis* used for comparison with sedimentary eDNA.
